# Macroecological Laws Can Naturally Arise from Chaotic Internal Species Dynamics

**DOI:** 10.1101/2025.10.05.680583

**Authors:** Konstantine Tchourine, Martin Carballo-Pacheco, Dennis Vitkup

## Abstract

Macroecological relationships that connect various statistical descriptors of long-term and short-term species dynamics represent some of the most general laws in ecology and biology. These macroecological laws have been observed across diverse ecologies of plants and animals, and more recently, also in microbiota. Yet it remains unclear why strikingly similar macroecological relationships often arise in very different biological communities and various environmental contexts. Here, we investigated whether chaotic internal dynamics in spatially heterogeneous communities could underlie multiple macroecological relationships. Our analyses reveal that very general constraints on species interactions and spatial migration parameters can simultaneously lead to multiple macroecological laws found in microbial ecosystems without requiring external sources of noise. Our study also identifies the mechanistic origins of many empirically observed macroecological relationships, such as Taylor’s law, anomalous abundance diffusion, the Laplace distribution of short-term abundance changes, and the distribution of species residence times. Overall, we demonstrate how macroecological laws can arise from interaction-driven chaotic dynamics and common ecological constraints, thereby providing a unifying explanation for their widespread prevalence in nature.

## Introduction

Over the past two decades, large-scale studies have demonstrated that the microbiota plays an important role in human biology and disease^1–4^. Analysis of the essential physiological functions played by animal microbiota requires an understanding of the complex ecosystem dynamics, which often involves hundreds of interacting microbial species^2,5^. As in many other ecological systems, substantial temporal and spatial variability are salient features of the animal microbiota^6–9^. Recently, to better characterize and understand the structure and dynamics of diverse microbial communities, researchers have increasingly adopted approaches from quantitative and statistical ecology^10–13^.

We and others have previously demonstrated that, despite a many orders of magnitude difference in the characteristic time and length scales, microbiota dynamics often follow multiple macroecological relationships that are strikingly similar to those previously described in diverse ecologies of plants and animals^7,14–16^. These macroecological scaling laws typically characterize ecological abundance distributions and correlations between species’ abundance and their temporal and spatial variability. Briefly, on short time scales, the daily bacterial abundance changes 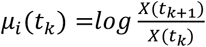, where *X*_*i*_(*tk*) is the relative abundance of microbial species *i* on day *k*, follow the Laplace distribution with the probability density 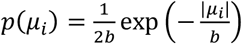, with a scaling parameter *b*. In contrast to the Gaussian distribution, the Laplace distribution implies a substantially higher probability of large-scale fluctuations in microbiota and other ecosystems^17,18^. Furthermore, the standard deviation of daily abundance changes, *μ*_*i*_, scales linearly with mean species’ daily abundances, indicating that the fluctuations of relative abundances are more pronounced for low-abundant microbial species, consistent with observations in other ecosystems^19^.

On longer time scales, the slow mean-squared temporal drift of log bacterial abundances, ⟨*δ*^2^(Δt)⟩, is well described by the equation of anomalous diffusion, ⟨*δ*^2^(Δt)⟩ ∝ Δ*t*^2*H*^, where Δ*t* is the time lag between observations, and *H* is the so-called Hurst exponent^20,21^. Notably, similar long-term anomalous diffusion of species abundances has been described in diverse animal communities^17,22^. The relationship between temporal means and variances of microbiota species abundances follows a power law, known in ecology as Taylor’s law^23–26^, 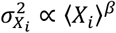, where the abundance variance, 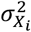, scales with mean abundance, ⟨*X*_*i*_⟩, with a positive exponent *β*, across species *i*. Another characteristic of long-term ecological dynamics, the distribution of species’ residence and return times, has also been established in various communities^17,27,28^. In microbiota, as in animal ecologies, these distributions follow a power law with an exponential cutoff tail, *p*(*t*) ∝ *t*^−*α*^*e*^−*λt*^. In addition to the macroecological relationships described above for microbiota dynamics^7^, several other scaling laws are known to characterize structural properties of multiple ecosystems. For example, the species abundance distribution often follows a scaling relationship, 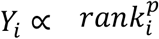, where *Y*_*i*_ is the mean abundance of a species *i, rank*_*i*_ is the abundance rank of species *i* compared to all other species, and *p* is the power law exponent^29–31^.

Despite their prevalence in nature, it remains unclear why so many macroecological relationships consistently arise across vastly different environmental contexts and ecosystems. This observation suggests that commonly observed macroecological laws are unlikely to result from ecosystem-specific interspecies interactions or particular patterns of environmental fluctuations. Instead, macroecological laws may originate from general properties of species interactions and constraints that are shared between ecosystems. In this study, we investigated whether chaotic internal species dynamics can simultaneously give rise to multiple macroecological laws. To this end, we used a generalized and spatially-resolved Lotka-Volterra model to reproduce experimentally observed microbiota dynamics. We then examined how general constraints on the distribution of key system parameters influence the observed macroecological relationships. Using optimized models, together with available experimental data, we derived mechanistic insights into the origins of multiple macroecological laws that characterize both the short- and long-term ecological dynamics. Overall, our analysis demonstrates how diverse macroecological laws can simultaneously emerge from complex interactions between species and common constraints likely shared among ecosystems.

## Results

### Optimizing the hyperparameters of the generalized Lotka-Volterra model

Many natural ecosystems, including microbiota, are not well-mixed communities^32–34^. Instead, they are typically characterized by substantial spatial heterogeneity^6,35–37^, with frequent exchange of species between sub-communities at different spatial locations^38,39^. Therefore, to investigate the origins of macroecological relationships, we used a spatially resolved generalized Lotka-Volterra (sgLV) ecosystem model. Recent studies have demonstrated that sgLV ecosystem models can produce large fluctuations in species abundances while also preserving species diversity^40–42^. These elegant studies followed earlier ecological analyses demonstrating that spatial heterogeneity is crucial for preventing substantial species loss and maintaining experimentally observed abundance fluctuations^38,43,44^.

The sgLV model we used includes multiple spatially separated locations (“islands”), which were all assumed to be internally well mixed (see Methods). Species interactions at each spatial location were approximated by generalized Lotka-Volterra dynamics, and species were also allowed to migrate between the spatial locations (Figure 1a, see Methods). The dynamics of abundance, *x*_*i,u*_, of each species *i* on island *u* were described by the differential equation:

**Figure 1:**
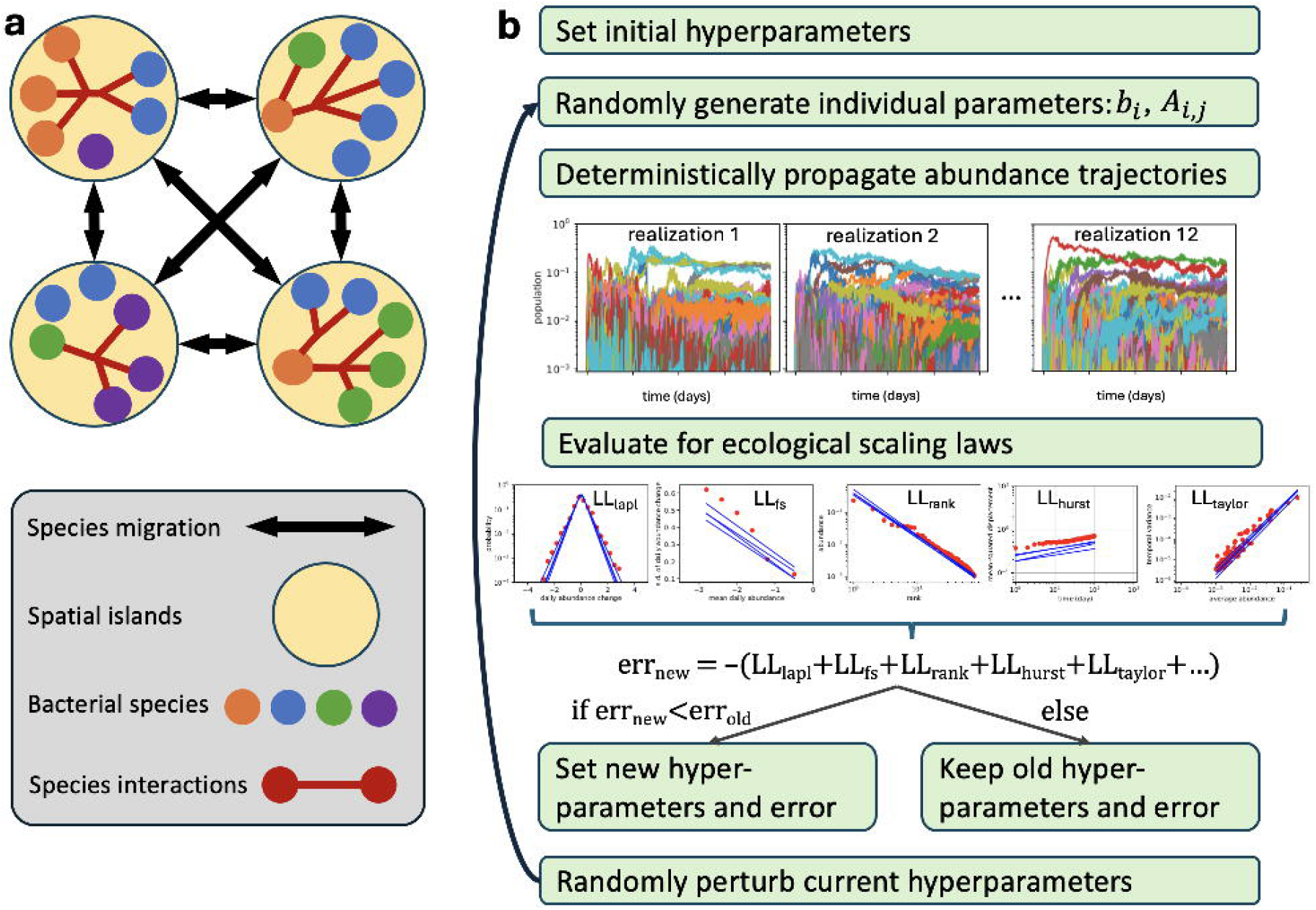
Model outline and optimization algorithm. **a**, Ecological model. The abundance of each bacterial species (colored dots) can change due to intrinsic growth, interactions with other species (red lines), and migration between islands (black arrows). **b**, Simulated annealing for hyperparameter optimization. In each iteration, the current hyperparameters (error = err_old_) are randomly perturbed to produce the updated hyperparameters. The updated hyperparameters are used to generate 12 realizations of individual species parameters (growth rates *b*_*i*_ and interactions *A*_*i,j*_), and abundance trajectories are simulated. Each realization is scored against ecological scaling laws (Laplace-distributed fluctuations, Taylor’s law, etc) by computing the log-likelihood (LL) of the simulated data relative to the four human time-series datasets in Ji et al. (2020)^7^. The realization-specific error is defined as the sum of negative log-likelihoods across scaling laws. The new total error (err_new_) is the average of the eight best realizations. Candidate hyperparameters that decrease the error are always accepted; otherwise, acceptance occurs with a probability that decreases with both simulated annealing step and error increase (err_new_ − err_old_.). This process is repeated until convergence.

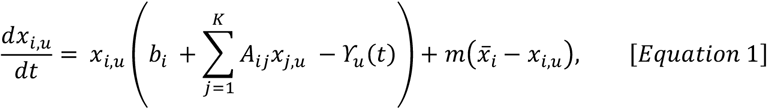

where the parameter *b*_*i*_ represents the intrinsic growth rate of species *i*, the parameters *A*_*ij*_ describe the interaction strength of species *j* with species *i, K* is the total number of species, *m* is the species migration rate between the spatial locations, and 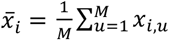 is the mean abundance of species *i* across all *M* spatial locations. We also introduced a carrying capacity parameter *ϒ*_*u*_(*t*) (see Methods), which ensures that the total abundance, 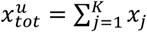, of all species at each spatial location *u* can fluctuate according to the logistic equation with a characteristic parameter *α*_*u*_ (see Methods)

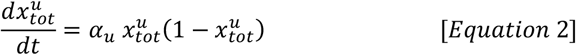

A fully specified sgLV model contains many hundreds of growth and interaction parameters. Therefore, to avoid model overfitting and to gain a mechanistic understanding of the origins of macroecological laws in natural ecosystems, we introduced a small set of adjustable hyperparameters. Rather than specifying the growth and interaction parameter values for each individual species and pair of species, these hyperparameters define the shapes and spreads of the parameter distributions across species. We then asked whether it is possible to optimize the hyperparameters to maximize the fraction of random parameter realizations –sampled from hyperparameter-constrained distributions – that produce models consistent with experimentally observed ecological dynamics. If this is indeed the case, it would suggest that certain constraints on the structure and strengths of interactions and growth rates across species can explain the frequent occurrence of macroecological laws in nature.

We first optimized the hyperparameters to fit five macroecological relationships that describe microbiota dynamics^7,16^, specifically: the power-law distribution of ranked species abundances, which represents an essential structural property of the simulated ecosystem (Figure 2a, Supp. Figure 1), the relationships characterizing the short-term microbiota dynamics, i.e., the Laplace distribution of species’ abundance fluctuations (Figure 2b) and the short-term fluctuation scaling with species’ daily abundances (Figure 2c), and the relationships characterizing the long-term microbiota dynamics, i.e., the Hurst anomalous diffusion of microbiota abundances (Figure 2d) and the scaling of the temporal abundance fluctuations with the average bacterial abundances (Taylor’s law, Figure 2e). To fit these macroecological laws, we optimized seven free hyperparameters: the mean and variance of species’ growth rates *b*_*i*_, the mean, variance, and symmetry of interactions *A*_*ij*_, the species migration rate between islands *m*, and the mean of total abundance response *α*_*u*_. To optimize the hyperparameters, we used a simulated annealing algorithm (see Methods, Figure 1b, Supp. Figure 2). Briefly, optimization runs were initialized from a random set of hyperparameters. Hyperparameter-constrained distributions were then used to generate multiple (n=12) sets of random parameter realizations for all species. Each parameter realization was then used to simulate 400-day-long abundance trajectories for all species in the ecosystem. At each optimization step, the five macroecological relationships calculated from the simulated dynamics were evaluated for goodness-of-fit to the experimentally observed macroecological scaling laws, which were derived based on one-year long microbiota trajectories^1,45^. Based on the fits, the new hyperparameters were either accepted or rejected, and iterations were continued until hyperparameter convergence was reached (see Methods, Supp. Figure 2).

**Figure 2.**
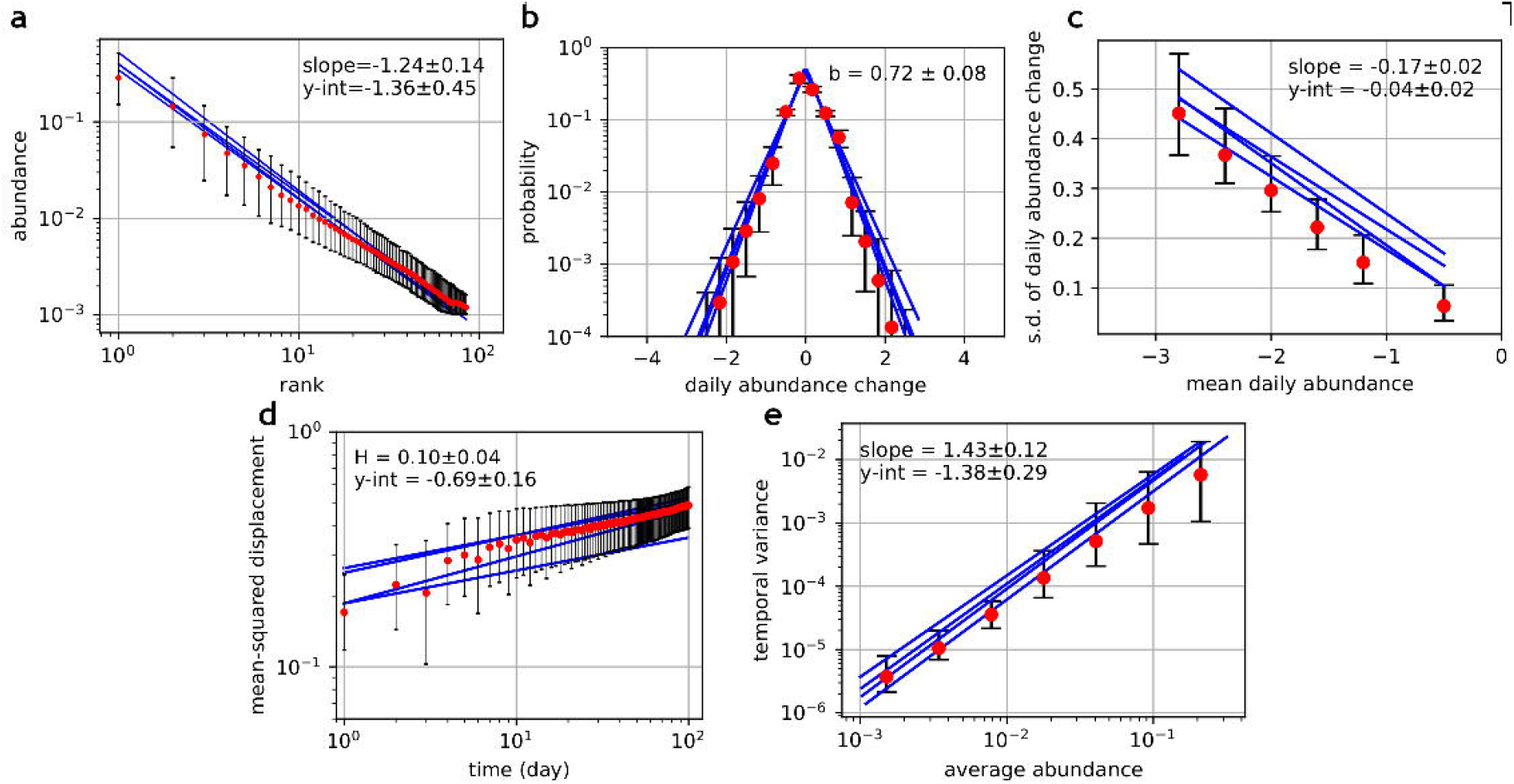
Simulated trajectories follow the optimized ecological scaling laws. Results from the best (lowest-error) 250 out of 500 random realizations of the optimized hyperparameters are shown. **a**, Mean relative species abundance as a function of the species’ rank among all species. **b**, Distribution of daily abundance changes. The value of the Laplace distribution exponent, *b*, calculated on the fit to the simulated data, is reported. **c**, Scaling of the standard deviation of daily abundance changes with mean daily abundances. **d**, Mean-squared displacement, ⟨*δ*^2^(Δ*t*)⟩, scales linearly on a log-log plot with time lag Δ*t*. The low Hurst exponent *H* = 0.10 ± 0.04 is characteristic of a subdiffusive process. **e**, Mean abundance of each species is linearly proportional to the variance of each species, evaluated across time. In all panels, red dots and error bars show median and 90% interquantile range across the 250 realizations, respectively, plotted within the experimentally-measured range, while the four blue lines represent the fits to the human data^1,45^. Scaling exponents represent means and standard deviations of exponents calculated in the 250 best realizations.

### The sgLV model accurately approximates multiple empirically observed distributions and macroecological laws

A substantial fraction of ecological models based on species interactions and growth parameters randomly sampled from the distributions specified by the optimized hyperparameters accurately approximated the five empirically observed macroecological laws used for fitting (Figure 2), with over 60% of random realizations having a smaller total log-likelihood error (see Methods) than the typical total error between pairs of human experimental profiles. Reassuringly, even without explicit optimization, the simulated microbiota trajectories reproduced several other experimentally observed ecological distributions. For example, long-term species dynamics can be characterized by the distributions of their residence and return times^7,17,27,28^. Without specific optimization to fit these distributions, our simulations accurately captured the corresponding scaling relationships observed in the experimental data (Figure 3a-b). The distribution of temporal abundance correlations between pairs of simulated species also closely matched the distribution observed in the experimental data (Figure 3c). Furthermore, the distribution of Hurst coefficients across individual species, characterizing the long-term anomalous diffusion of their log abundances, accurately approximated the experimental distribution (Figure 3d, Supp. Figure 3), with over 80% of the simulated species showing significantly positive (Bonferroni-corrected P<0.01) Hurst exponents (median individual species Hurst exponent *H* = 0.07).

**Figure 3.**
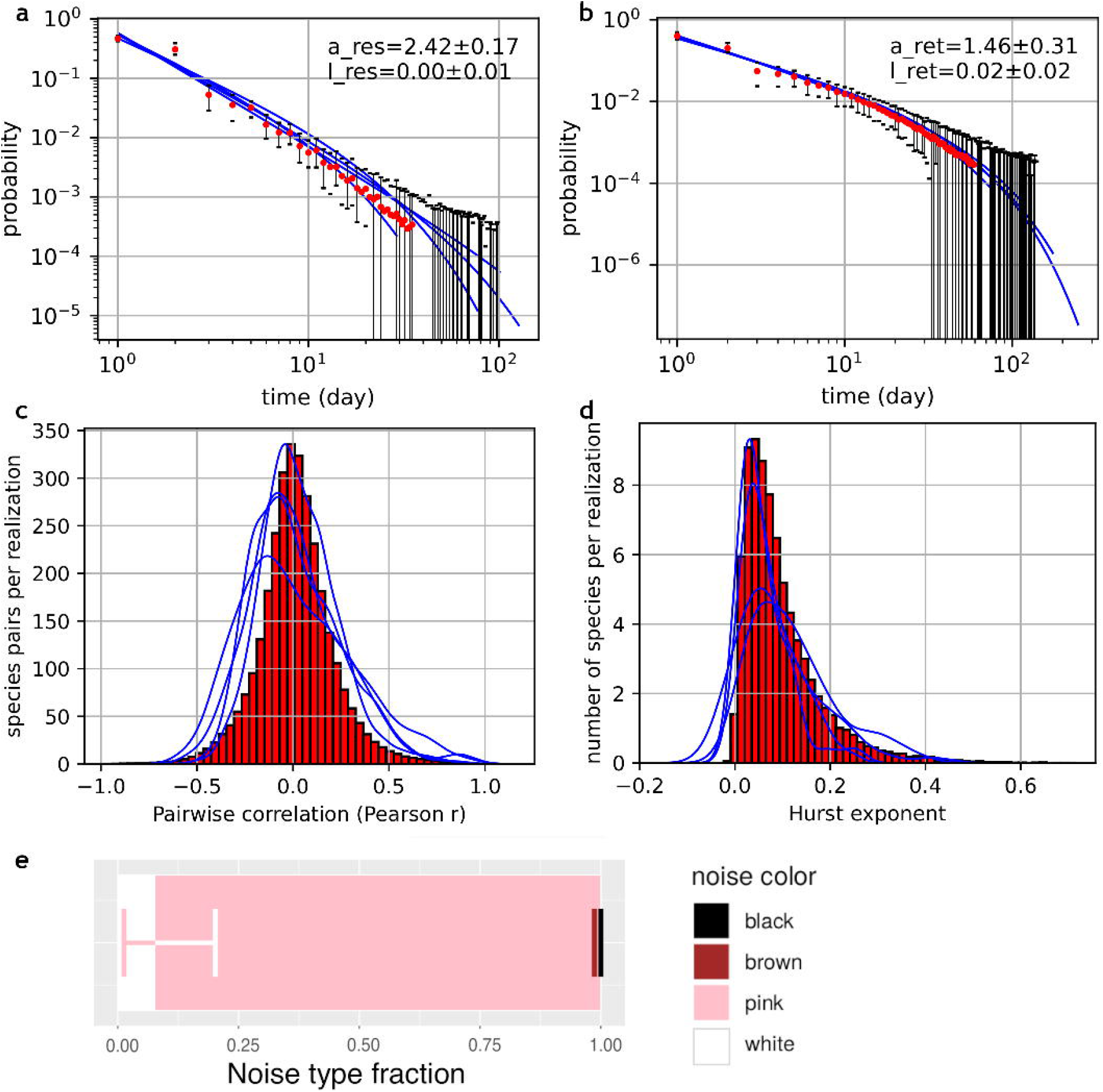
Simulated trajectories follow unoptimized ecological scaling laws. **a**-**b**, Distributions of residence (**a**) and return (**b**) times follow a power law with an exponential cutoff. Shown on the plots are means and standard deviations of *α*_*res*_ and *α*_*ret*_, the coefficients of power law fits to residence and return times, respectively, with the corresponding exponential cutoff exponents *λ*_*res*_ and *λ*_*ret*_, calculated across hyperparameter realizations. Red dots and error bars show median and 90% interquantile range, respectively. **c**, Distribution of pairwise species correlations across simulated species. **d**, Distribution of Hurst law exponent *H*, calculated individually in each species. In **a-d**, the blue curves represent the fits to the experimentally-measured human data^1,45^, and red dots and bars show simulated data in the best 250 (out of 500) hyperparameter realizations. **e**, Noise color composition showing the median fraction of high-abundance species with a noise spectrum in each category, across the best 68 (out of 96) optimal hyperparameter realizations. Error bars denote the 68% interquantile range of each noise color.

The temporal structure of species abundance trajectories can be analyzed using their temporal autocorrelation. Based on the Fourier transform of the autocorrelation function, the temporal structure of the abundance trajectories can be described by predominant color types, which characterize the relationship between spectral density and temporal frequency^11,14^. Fourier decomposition analysis revealed that both experimental and simulated trajectories were dominated by pink noise, with spectral density inversely proportional to frequency (Figure 3e, Supp. Figure 4). Indeed, pink noise is frequently observed in diverse natural processes that exhibit long-term temporal memory^11,46,47^.

Overall, our results demonstrate that ecological species dynamics simulated using the optimized hyperparameters can simultaneously capture various key features of experimental data, including multiple macroecological relationships and associated statistical distributions observed in gut microbiota and in many other ecosystems. Interestingly, accurate fits to the experimental data were obtained without the addition of external noise sources or specific environmental perturbations. This demonstrates that the empirically observed macroecological relationships can arise mechanistically from internal chaotic species dynamics.

### Insights into the mechanistic nature of macroecological laws

We next used simulations with optimized hyperparameters to gain mechanistic insights into the origins of empirically observed macroecological relationships. First, we investigated which general properties of model parameter distributions were crucial for reproducing macroecological laws. To that end, we performed a hyperparameter sensitivity analysis (Supp. Figures 5-6) using the optimized set of hyperparameters as a starting point (see Methods). This analysis showed that macroecological laws were most accurately approximated in simulations when interactions between species were dominated by competitive (mutually detrimental) and parasitic (predator-prey) interactions, consistent with observations in real ecosystems^10,48,49^. We also found that other properties of the species interaction matrix, such as the mean interaction strength, affected both the short-term and long-term dynamics of the simulated system (see below), with optimal hyperparameter values representing a balance (Supp. Figure 5) between rapid fluctuations and the long-term stability of the ecosystem^40,50–52^. In contrast to properties of the species interactions matrix, hyperparameter optimization and sensitivity analysis showed that macroecological laws were not sensitive to the mean and variance of the species growth rate hyperparameters (Supp. Figure 2 and Supp. Figure 6). This indicates that the macroecological relationships depend more on the properties of species interactions between species within the ecosystem than on growth rate parameters.

We next investigated the mechanistic origins of individual macroecological laws. We and others have reported that short-term abundance changes in microbiota, as well as in other ecosystems, follow the Laplace distribution^7,17,19^, but why this particular distribution is commonly observed in ecology is not well understood^53–56^. Previously, we found that the overall Laplace distribution is unlikely to arise as a mixture of Gaussian distributions representing daily abundance changes of individual OTUs^22,57^, because abundance changes calculated for each OTU individually also followed the Laplace distribution^7^. To further examine the origin of the distribution of species daily abundance changes, we investigated how it depends on the length of the time window over which it is calculated. Notably, we found that the abundance changes for individual species, both in our simulations and in experimental data, followed a Gaussian distribution on shorter time scales, but increasingly approximated the Laplace distribution as the time window over which the distribution was calculated increased (Figure 4a). To assess how the Laplace distributions are formed, we examined how the means and the variances of abundance change distributions observed over 30-day time windows varied across the entire 320-day simulation trajectory. We found that individual species with greater variability in the variance of the 30-day abundance change distributions were more likely to exhibit a Laplace distribution of abundance changes across the entire trajectory (Figure 4b). In contrast, greater variability in the means of 30-day abundance change distributions was not generally associated with more Laplace-like distributions (Figure 4c). This suggests that the abundance change of each species behaves as a Gaussian process on the scale of a month, consistent with random multiplicative processes^58,59^, but fluctuations in the abundance variance across months result in a mixture of Gaussian distributions^22,57^, giving rise to the Laplace distribution of daily abundance changes over longer time periods.

**Figure 4.**
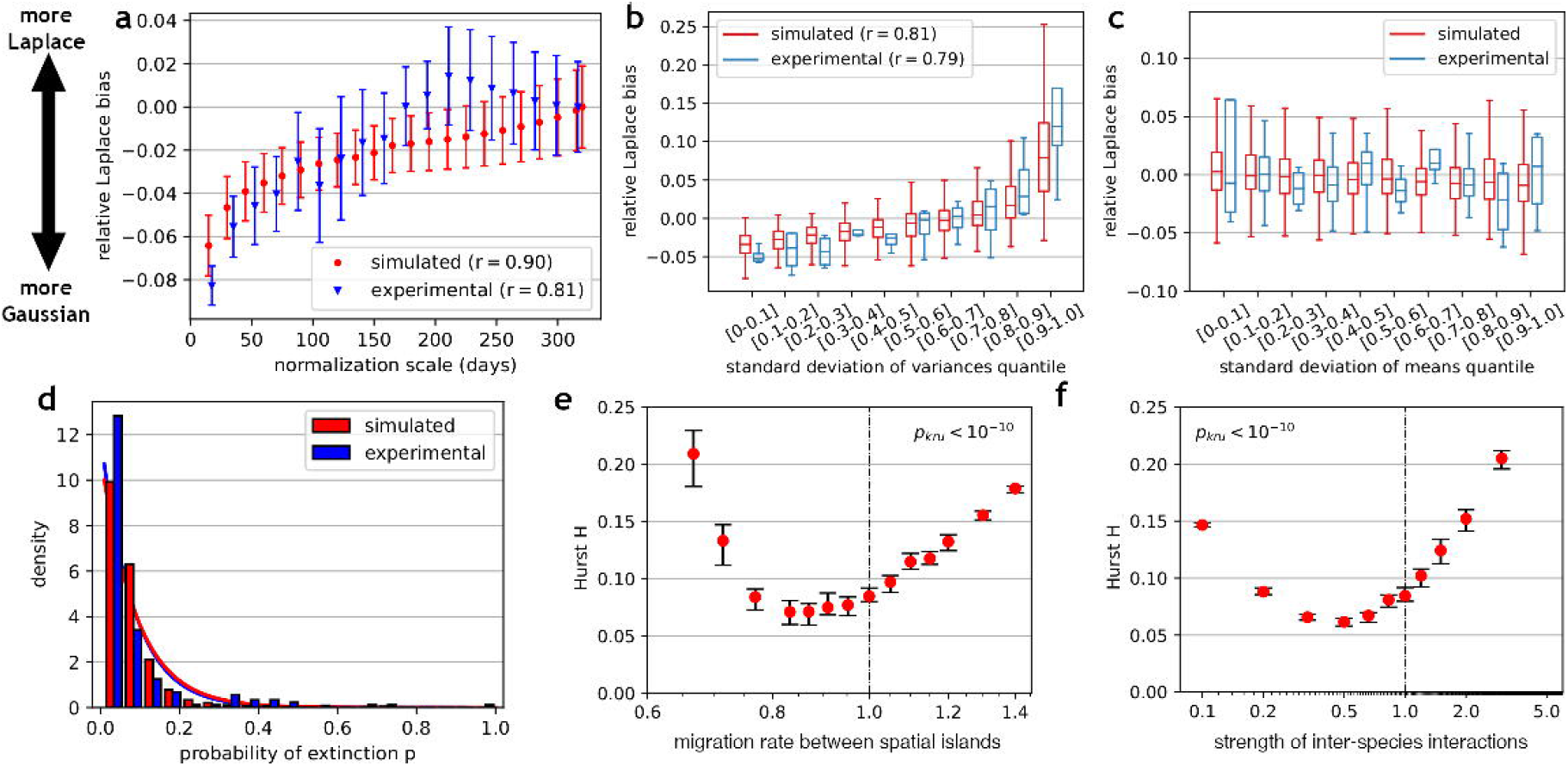
Determinants of individual scaling laws. **a**, Relative Laplace bias (see Methods) of the distribution of daily abundance changes as a function of the length of intervals across which individual species abundance changes are normalized to the same variance. Shapes and error bars represent the means and variances of relative Laplace bias across the best 250 (out of 500) random realizations of the optimal hyperparameters. **b**, Box plots of relative Laplace bias as a function of the variability of species’ 30-day variances. For each species, the variance of daily abundance changes was calculated within every 30-day window, and the across-window variance of these values was used as a measure of “variance fluctuation”. Species were grouped into bins by this quantity, and relative Laplace bias was computed from the pooled daily abundance changes within each bin. The average correlation coefficient r across 250 realizations is reported. **c**, Same as in **b**, but species are binned by the variability of their 30-day means rather than their variances. **d**, The distribution of the rate parameters of individual species residence times, across simulated species (red) and experimentally measured^1,45^ OTUs (blue), approximately follows an exponential distribution (line fits). **e-f**, Hyperparameter sensitivity analysis shows the dependency of the Hurst exponent on the inter-island migration rate (**e**) and the mean inter-species interaction strength (**f**). Red dots and error bars show the median and 95% confidence interval on the scaling exponent median, calculated using a non-parametric bootstrap test across 300 random realizations of each hyperparameter. P-values from the Kruskal-Wallis test are reported.

To examine the origin of the power law distributions with exponential cutoffs (Figure 3a) that describe species residence times, we first calculated residence time distributions for individual species. Exponentially distributed extinction and lifespan times have been observed for many species in ecology^17,27,28^. Consistent with these observations, when we applied the Akaike Information Criterion to compare exponential and power law distribution goodness of fits for residence times of individual species, we found that at least two-thirds of individual microbial species had residence times that were more consistent with the exponential distribution, both in simulations (Supp. Figure 7a) and human gut data (Supp. Figure 7b; see also Tchourine *et al*. (2021)^55^). We then asked how species combine to produce the observed power law distributions with exponential tails. Interestingly, we found that combining residence times for only the geometrically distributed species, as well as for only the power law distributed species, yielded a similar distribution to that observed across all species (Supp. Figure 7c-f). To examine how the exponentially distributed residence times of individual species combine into a power-law distribution across all species, we calculated the distribution of the rate parameter of the exponential fit for each species. Notably, we found that the rate parameter follows the exponential distribution, both in human gut data and in our simulations (Figure 4d). This suggests that when combined across species, the power law distributions with exponential tails of residence times can arise as a combination of power-law distributed variables or as a combination of exponentially distributed variables with a broad distribution of the exponential rate parameter.

We next examined the factors that influence the long-term anomalous Hurst diffusion of microbial abundances. Our hyperparameter sensitivity analysis showed that the Hurst exponent was particularly sensitive to the migration rate (Figure 4e) and to the mean interaction strength between bacteria (Figure 4f). We found that the degree to which spatial separation is maintained through migration between islands strongly influences the global long-term drift of microbial abundances. When spatial islands are largely isolated, i.e., at low migration rates, rapid species extinction on individual islands is not counterbalanced by an influx of new species, resulting in greater overall species abundance drift (high H values). Conversely, when islands are connected too strongly through species migration, the entire ecosystem resembles a single well-mixed system, again leading to relatively rapid abundance diffusion. With the right balance between island isolation and homogenization through species migrations, the anomalous Hurst diffusion approximates the one experimentally observed in microbiota (Figure 2d).

Additionally, analysis of random realizations of the interaction matrix and the resulting abundance trajectories allowed us to investigate emergent ecosystem properties that were not explicitly controlled by the hyperparameters. For example, the strength and density of interactions of the species, which vary across different parameter realizations generated from the same hyperparameters, may influence the long-term dynamics of the system^50,52^. We quantified the strength of species interactions using the community matrix, a commonly used measure of ecosystem dynamics^60,61^. Specifically, we approximated the community matrix by multiplying each interaction coefficient *A*_*i,j*_ by the mean abundance of the affected species 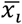, consistent with the definition of the community matrix in gLV models 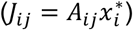^62^. This reflects the total influence exerted on population *i* by its interacting populations *j*. We found that Taylor’s law slope was significantly associated with the sum of the community matrix for the lowest 50% of species by abundance (Figure 5a, Spearman *r* = −0.34, *p* < 10^−7^, *n* = 250), while the relationship with the sum of the community matrix for the highest-abundance 50% of the species was not significant (*p* > 0.01). Stronger influence of the low-abundance species on one another increased their variability (Figure 5b, Spearman *r* = 0.57, *p* < 10^−22^), which in turn decreased Taylor’s law slope (Figure 5c, Spearman *r* = −0.38, *p* < 10^−9^). Overall, this analysis demonstrated that Taylor’s law slope depends on the total strength of the community matrix of low-abundance species, which may partially explain the variability of Taylor’s law across diverse ecosystems^7,12,63,64^.

**Figure 5.**
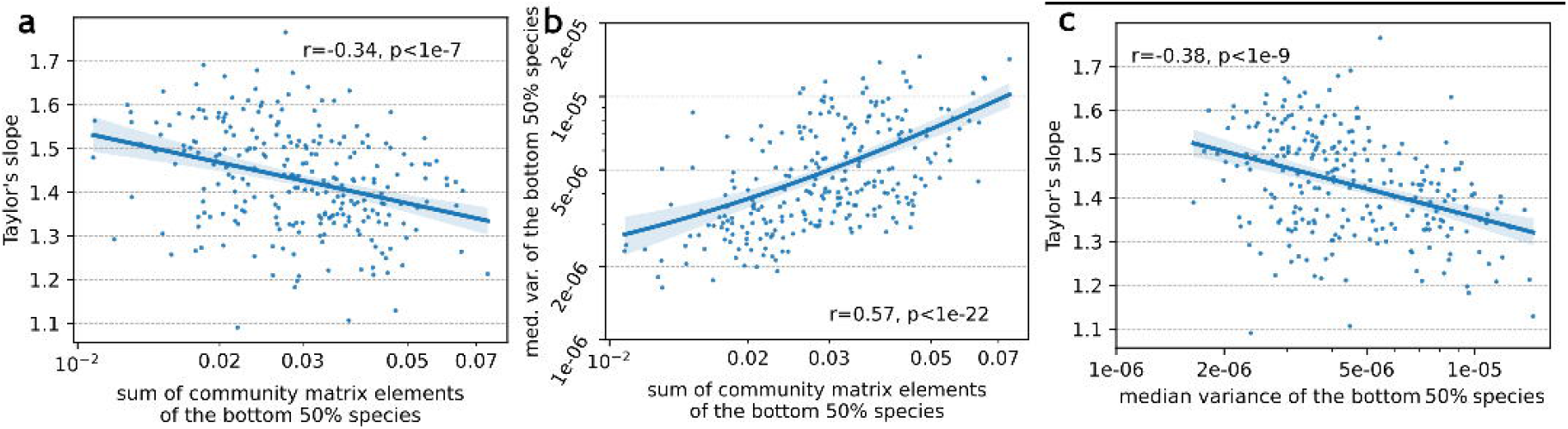
The community matrix is associated with the slope of the simulated ecosystem’s Taylor’s law. **a**, Taylor’s law slope varies with the sum of the community matrix values across the 50% lowest-abundance species. **b**, The median variance of the 50% lowest-abundance species is largely determined by the total strength of the community matrix elements for those species. **c**, Taylor’s law slope varies with the median variance of the 50% lowest-abundance species. On all panels, each dot represents a hyperparameter realization. The best 250 (out of 500) random realizations of the optimal hyperparameters are shown. Spearman correlation statistics are reported, and the blue line shows the log-linear fit to the data.

## Discussion

The field of macroecology has quantified multiple statistical patterns and dependencies exhibited by ecological communities ranging from animals and plants to microbiota^7,15,17,19,22,24,27,28^. Notably, similar macroecological laws are observed across ecosystems exposed to diverse environmental perturbations, as well as in relatively stable conditions, such as the mouse gut microbiota maintained on a constant diet^7,65^. This suggests that common functional and structural principles related to internal species dynamics may underlie these frequently observed macroecological laws. The pioneering work of Robert May demonstrated that internal species interactions, even without external environmental noise, can lead to complex and chaotic population dynamics^66^. In addition, various spatially resolved models have been developed in ecology to study the dynamic properties of ecological systems^38,43,44^.

In this study, using a spatially-resolved generalized Lotka-Volterra model, we demonstrate that commonly observed macroecological laws can naturally and simultaneously arise from internal dynamics in complex ecosystems, provided that interaction types and their strengths remain within a certain broad range. After we described multiple macroecological laws in microbiota^7,16^, several models attempted to capture various macroecological relationships in microbial ecology^14,54,56,67–75^. We note that the model presented here has several distinct and important features. First, our model reproduces all of the macroecological scaling laws characterized in the gut microbiota by Ji et al. (2020)^7^. In particular, it accurately captures the Hurst power law that describes the anomalous long-term diffusion both of the whole ecosystem and of individual species. Second, our model does not require stochastic perturbations or environmental noise to reproduce the experimentally observed macroecological laws.

Third, our model does not require defining and fitting individual carrying capacities for each species. Instead of imposing preset carrying capacities on individual species^14,68,76^, we implemented a flexible total load capacity on the entire ecosystem. This resulted in emergent species carrying capacities with a power-law rank-abundance scaling that closely approximates experimental observations.

We demonstrate in the paper that the parameters (exponents) of individual macroecological scaling laws depend on the interaction strength between species across time and spatial locations. Specifically, we found that a sparse matrix of relatively weak interactions was sufficient to reproduce the high short-term fluctuations observed in microbial communities. Additionally, weak coupling between spatial islands supported the maintenance of high levels of diversity over long periods of time. The average interaction strength was associated with the Laplace exponent, suggesting that short-term fluctuations may result from local interspecies interactions. The Hurst exponent was also sensitive to the interaction strength and spatial migration rate, suggesting that long-term drift may depend on the spatial dynamics of the ecosystem. Interestingly, we found that Taylor’s law slopes, which have been previously investigated using various mathematical models incorporating spatial structure^77^, interactions^78^, resource fluctuations^54^, and demographic manipulations^74^, were associated with the strength of the emergent community matrix, with stronger interactions resulting in higher abundance variance of low-abundance species and lower Taylor’s law slopes.

This work contributes to the growing understanding of how ecological interactions can produce stable but rapidly fluctuating ecosystems^40,42,54,67,71^, and we demonstrate that macroecological laws can arise directly from internal species dynamics without specific environmental perturbations. This may explain the prevalence of these laws across diverse natural ecosystems with vastly different environmental conditions. Our model, together with other recent models, lays the groundwork for future studies of how macroecological laws may be affected by new species colonization and abrupt environmental changes. The rapidly developing methods for spatial sequencing of microbiota can be used to further constrain these models with experimental data. These insights will be essential both for understanding the human microbiota and its effects on health and disease, as well as for predicting dynamic changes in many natural ecosystems.

## Acknowledgements

This work was supported by grants from the National Institutes of Health (grant numbers R35GM131884 and R01DK118044).

