## Supplementary Figures for "Macroecological Laws Can Naturally Arise from Chaotic Internal Species Dynamics"

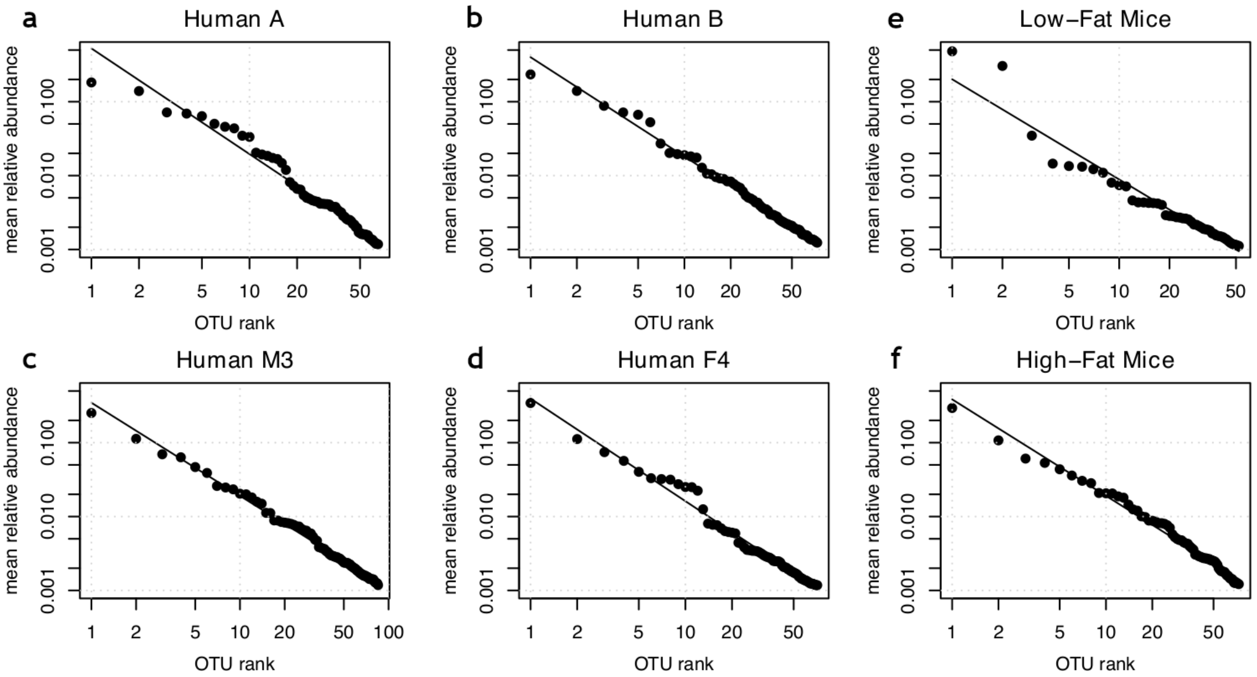


**Supplementary Figure 1. Rank-abundance scaling in experimentally-measured bacterial abundance data from Ji et al. (2020)[7]. a-d**, Rank-abundance scaling in four human datasets[1,45], shown on a log-log plot. **e-f**, The six mouse datasets[65] are grouped into two categories based on diet: low-fat plant-polysaccharide (LFPP) and high-fat high-sugar (HFHS) diets.

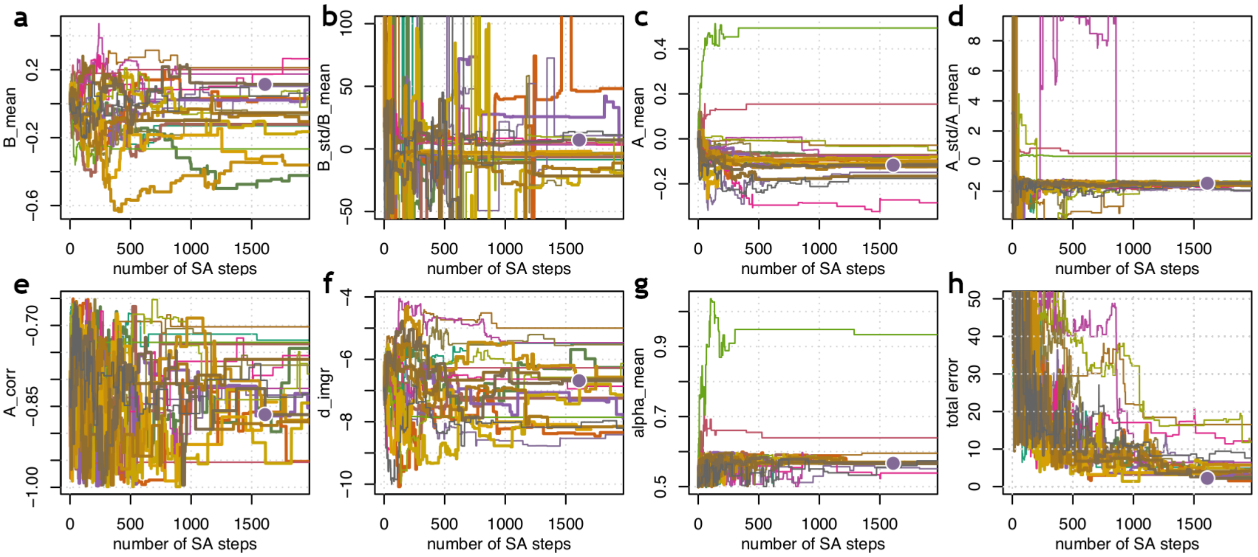


**Supplementary Figure 2. Hyperparameter convergence in simulated annealing optimization. a**-**g**, Values of the fitted hyperparameters as a function of the number of simulated annealing (SA) steps. **a**, Mean bacterial growth rate, $mean(b_i)$, **b**, Ratio of standard deviation of growth rate to mean growth rate, $\frac{std(b_i)}{mean(b_i)}$, **c**, Mean species interaction strength, $mean(A_{ij})$, **d**, Ratio of standard deviation of species interactions to mean species interaction, $\frac{std(A_{ij})}{mean(A_{ij})}$, **e**, Interaction symmetry, $cor(A_{ij}, A_{ji})$, **f**, inter-island migration rate $\text{d\_imgr} = \log(m)$ (see Equation 1 in the main text), **g**, Mean of total load response $\alpha_k$. **h**, Total error as a function of the number of simulated annealing steps. On all panels, the 20 individual lines represent independent optimization runs, with bolded lines representing 10 runs with the lowest minimum total error values. Purple dots show the optimal hyperparameter set used for further analyses throughout this paper.

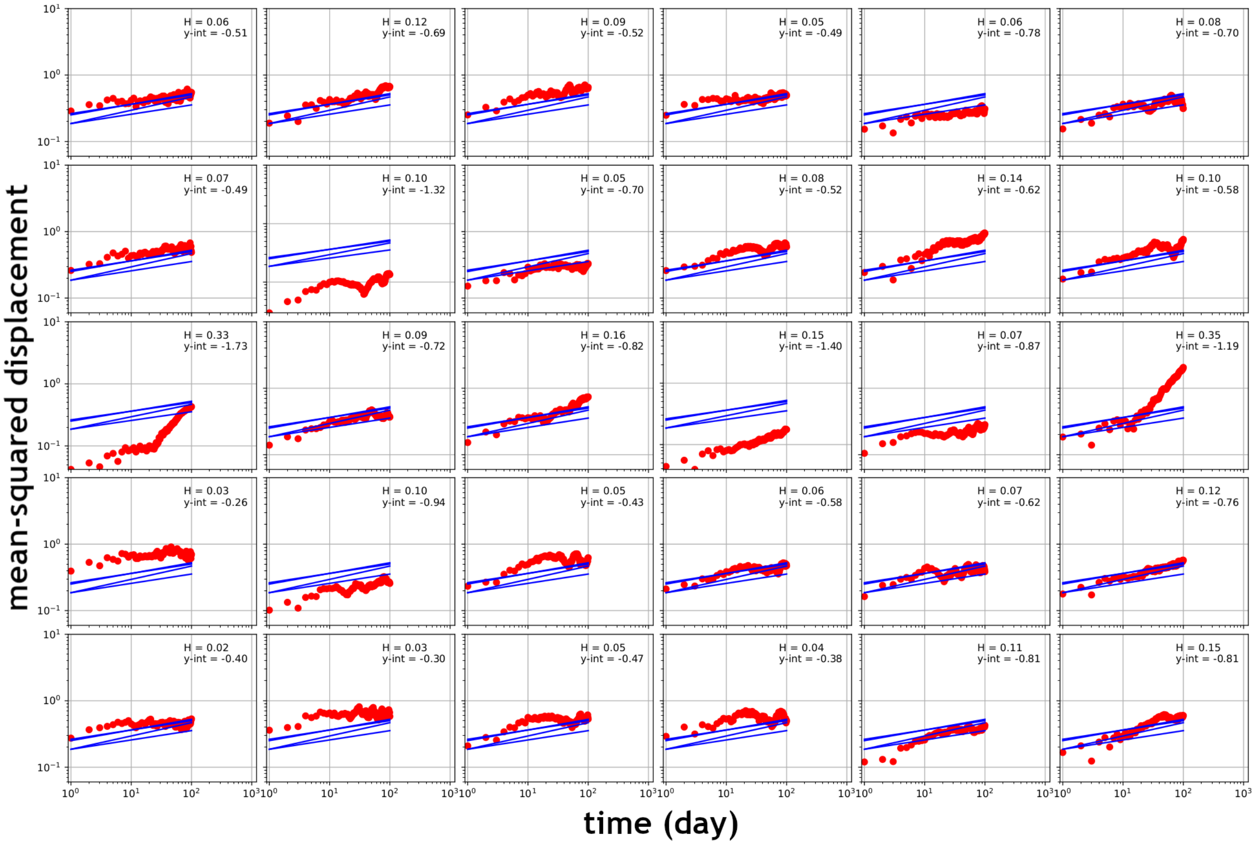


**Supplementary Figure 3. Long-term dynamics on an individual-species level.** Mean-squared displacement as a function of time lag $\Delta t$, calculated on each species individually, and displayed for the 30 most highly abundant species (individual panels). Displacements were calculated on data simulated using the optimized hyperparameters.

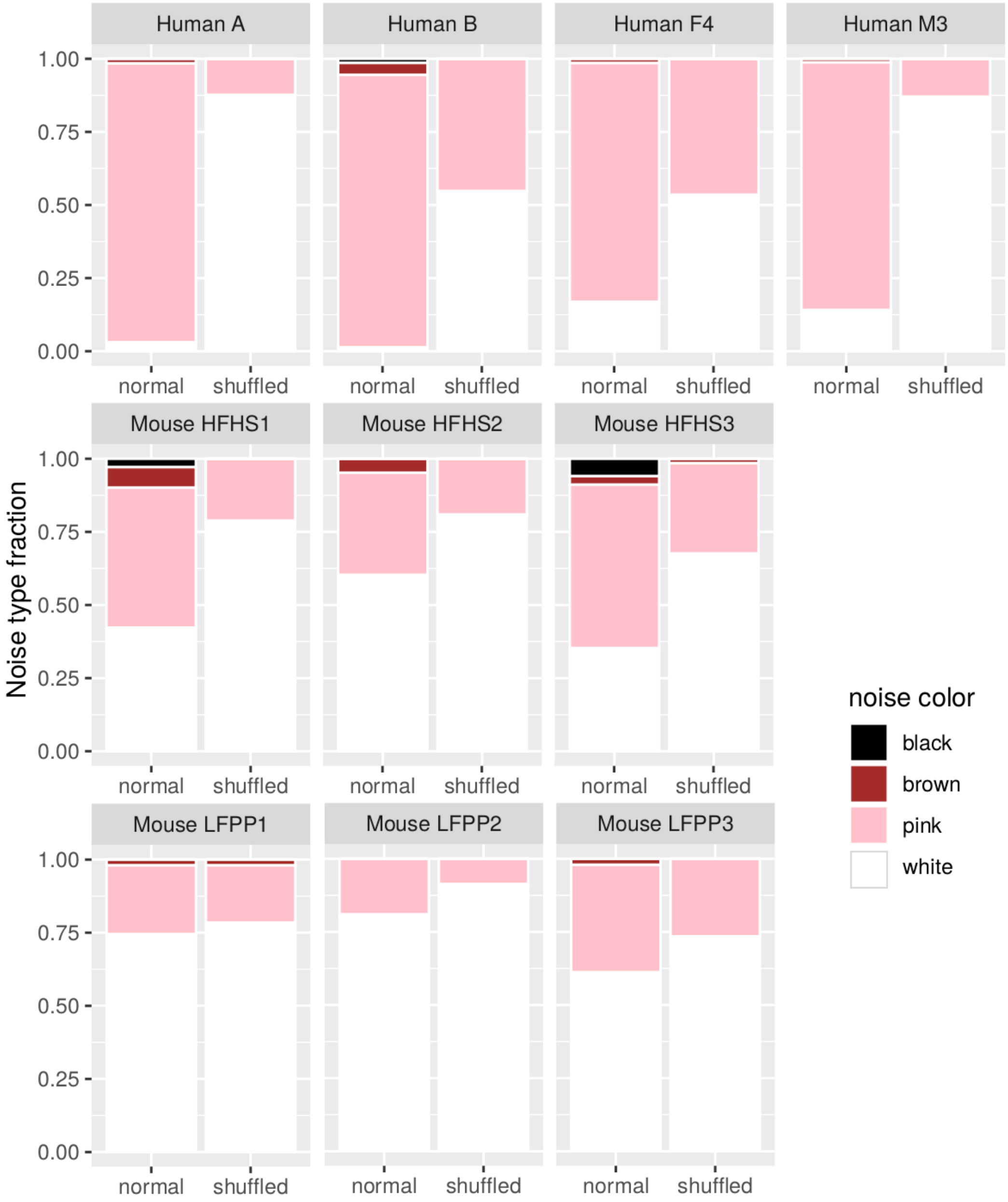


**Supplementary Figure 4. Noise decomposition profiles for human and mouse datasets.** For each of the human[1,45] and mouse[65] datasets in Ji *et al.* (2020)[7], spectral decomposition was performed for each OTU. The fraction of each noise color is represented by the height of the bar with the corresponding color. Noise decomposition and noise color assignments were performed as in Faust *et al*. (2018)[11] (see Methods).

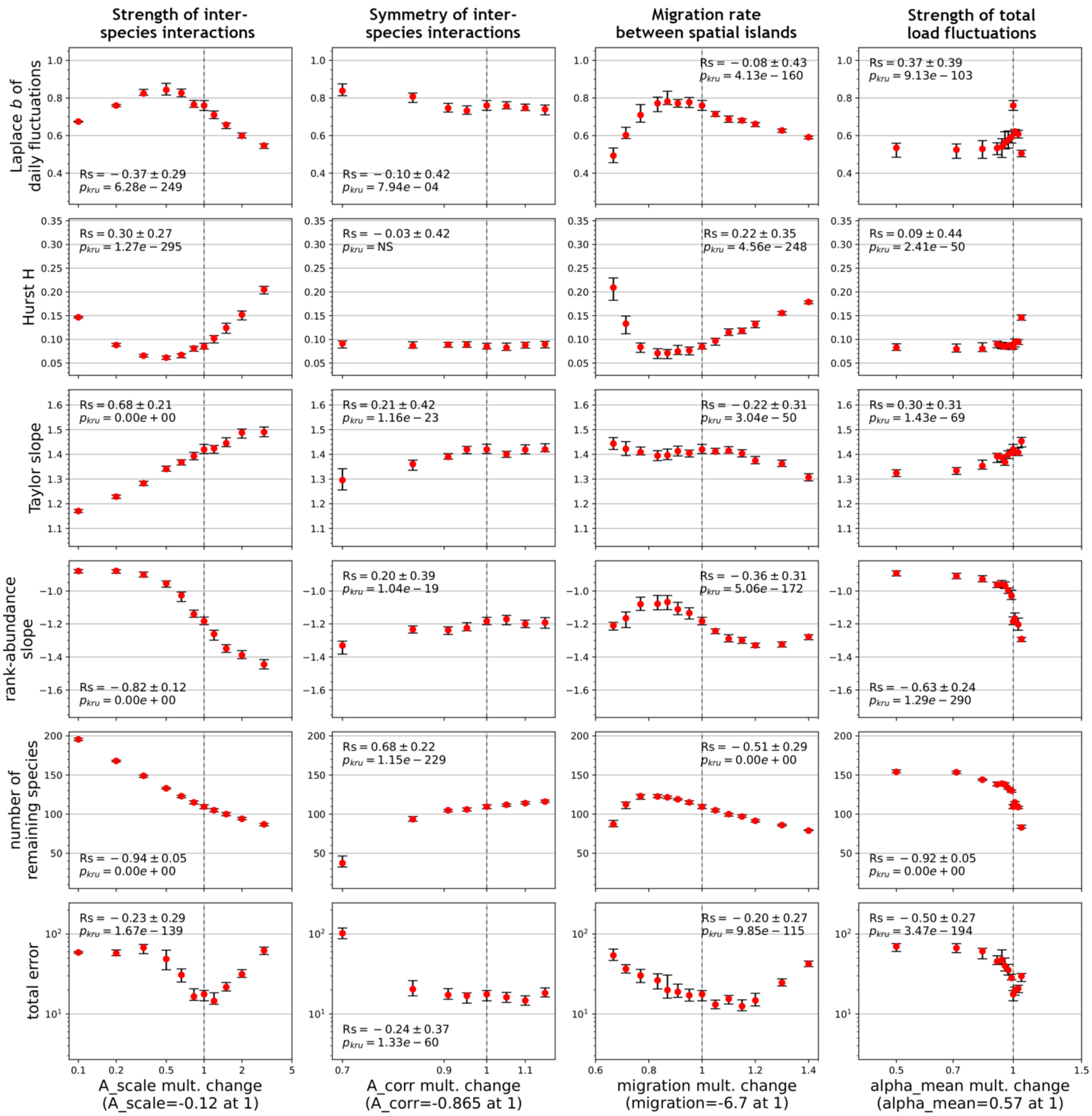


**Supplementary Figure 5. Hyperparameter sensitivity analysis.** Each panel shows the sensitivity of the corresponding scaling law statistic (rows) to variation in the corresponding hyperparameter (columns). The five scaling law statistics shown are, from top to bottom: the scale parameter $b$ of the Laplace distribution of daily abundance changes, the long-term drift coefficient $H$, the slope of the power law scaling between species means and variances (Taylor's law), the slope of the rank-abundance scaling, and the number of species with non-zero abundances in the last 100 days of the simulation. The lowest row shows the total log-likelihood error calculated from all fitted laws. The four hyperparameters shown are the mean species interaction strength $mean(A_{ij})$, the correlation strength $A_{corr}$ between $A_{ij}$ and $A_{ji}$, the migration parameter $m$, and the mean of the total abundance fluctuation parameter, $mean(\alpha_k)$. The x-axis on each subplot shows the multiplicative change in each hyperparameter compared to its optimal value (dashed line). Red dots represent the median of the scaling law statistic, with error bars denoting the 95% confidence interval of the median, calculated across the best 200 out of 300 realizations of the varied hyperparameter (x-axis), while keeping all other hyperparameters constant. P-values from the Kruskal-Wallis test and Spearman correlation coefficients are reported.

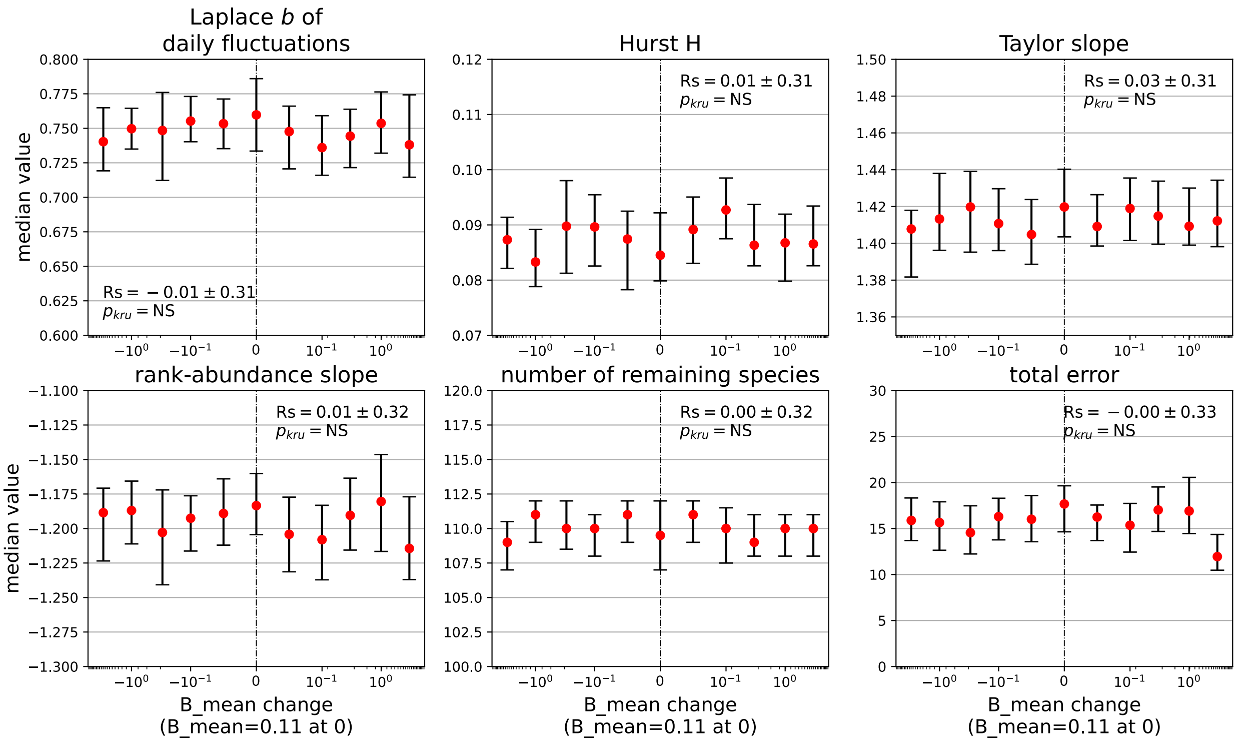


**Supplementary Figure 6. Sensitivity to the mean species growth rate.** Each panel shows the sensitivity of the corresponding scaling law statistic to variation in B_mean, the mean of growth rate $b_i$. The five scaling law statistics shown are: the scale parameter $b$ of the Laplace distribution of daily abundance changes, the long-term drift coefficient $H$, the slope of the power law scaling between species means and variances (Taylor's law), the slope of the rank-abundance scaling, and the number of species with non-zero abundances in the last 100 days of the simulation. The last panel shows the total error calculated from all fitted laws. Red dots represent the median of the scaling law statistic, with error bars denoting the 95% confidence interval of the median, calculated across the best 200 out of 300 realizations of the varied hyperparameter (x-axis, linear difference from optimal value), while keeping all other hyperparameters constant. P-values from the Kruskal-Wallis test and Spearman correlation coefficients are reported.

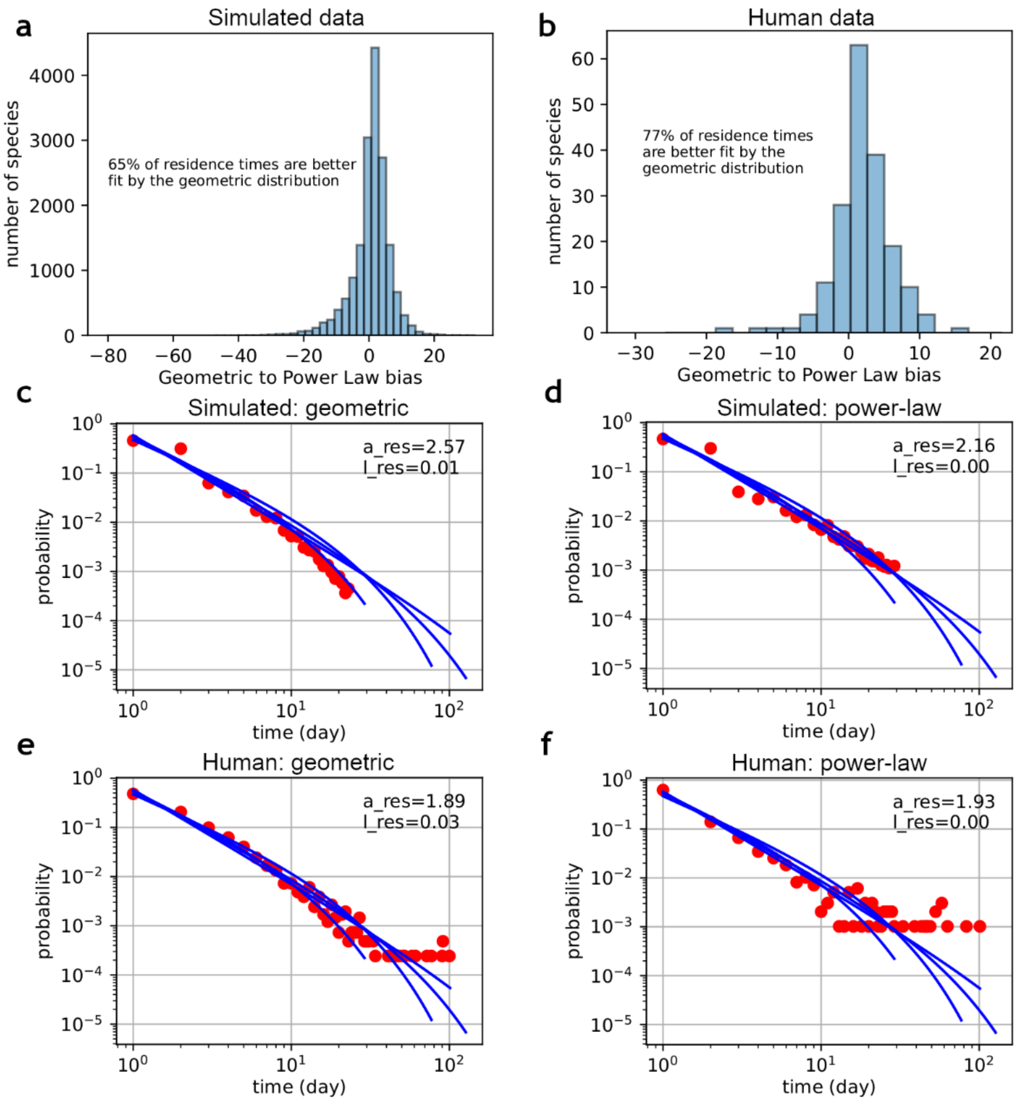


**Supplementary Figure 7. Residence times scaling arises as a combination of exponential and power law distributions for individual species. a-b,** The difference between the exponential (geometric) distribution and power law distribution fits to the residence times for each species for simulated trajectories (**a**) and experimentally-measured human datasets[1,45] (**b**). **c-f**, Distributions of residence times for simulated (**c**-**d**) or experimental (**e**-**f**) data in species with individual residence profiles better fit by exponential/geometric (**c**, **e**) or power law (**d**, **f**) distributions. Blue curves show fits to residence times of all OTUs in the four human datasets[7].
